# Retractions and rewards in science: An open question for reviewers and funders

**DOI:** 10.1101/2022.05.31.494225

**Authors:** Mariana D Ribeiro, M Kalichman, Sonia MR Vasconcelos

## Abstract

In recent years, the changing landscape for the conduct and assessment of research and of researchers has increased scrutiny of the reward systems of science. In this context, correcting the research record, including retractions, has gained attention and space in the publication system. One question is the possible influence of retractions on the careers of scientists. It might be assessed, for example, through citation patterns or productivity rates for authors who have had one or more retractions. This is an emerging issue today, with growing discussions in the research community about impact. We have explored the influence of retractions on grant review criteria. Here, we present results of a qualitative study exploring the views of a group of six representatives of funding agencies from different countries and of a follow-up survey of 224 reviewers in the US. These reviewers have served on panels for the National Science Foundation, the National Institutes of Health, and/or a few other agencies. We collected their perceptions about the influence of self-correction of the literature and of retractions on grant decisions. Our results suggest that correcting the research record, for honest error or misconduct, is perceived as an important mechanism to strengthen the reliability of science, among most respondents. However, retractions and self-correcting the literature at large are not factors influencing grant review, and dealing with retractions in reviewing grants is an open question for funders.

## Retractions and their influence on science indicators

Retractions and other types of corrections have gradually influenced the *modus operandi* of publishing science, but the extent to which they have affected or will affect the dynamics of doing, communicating and assessing scientific research is an open question. For example, it is not clear whether retractions influence indicators of research performance. Recently, it was shown that the number of retractions was higher for a group of highly productive countries, although a correlation between number of retractions and scientific productivity could not be established (Ribeiro & Vasconcelos, 2018). Concerning the impact of retractions on the career/performance of scientists, a growing number of scholars have shown that this impact may be assessed, for example, through the lens of citations (Furman et al. 2012; Lu et al. 2013). Lu et al (2013) found that citations dropped for authors of retracted publications. Specifically, “a single retraction triggers citation losses through an author’s prior body of work” and that “compared to closely-matched control papers, citations fall by an average of 6.9% per year for each prior publication…”. However, Lu and colleagues (2013) report that “citation losses among prior work disappear when authors self-report the error.” Mongeon and Larivière (2016) found that coauthors of a retracted publication, especially for misconduct, experienced a decrease in their individual productivity. Azoulay et al (2017) showed that “eminent scientists are more harshly penalized than their less distinguished peers in the wake of a retraction, but only in cases involving fraud or misconduct”.

## Retractions and the reward systems of science

These recent findings offer insight into the possible influence retractions might have on the reward systems of science, going beyond citations and productivity. Receiving less attention is the role that the correction of the literature has on the dynamics of research grant review. To our knowledge, criteria for evaluating grant proposals in major funding agencies rarely, if ever, include retractions or contributions of applicants to strengthening the reliability of science – for example, self-correcting the research record. In mainstream review criteria, self-corrections of the research record do not appear to count in funding decisions. On the other hand, retractions due to misconduct may lead to debarment from funding for a period of time. This is the case for major funders in the US, Europe and Latin America, which include the National Institutes of Health (NIH), the National Science Foundation (NSF) (Galbraith 2017), the German Research Foundation (DFG) (DFG 2018), and São Paulo Research Foundation (FAPESP) (FAPESP 2012). However, one question is whether and how one or more retractions – irrespective of the reason - in an applicant’s research record would influence funding decisions. How would reviewers’ assessments of new grant applications change, if at all, if it were known that the applicant was an author on a publication retracted for misconduct? In light of an increasing focus on researchers *getting it right* rather than *being right* or *being first* in research (Ebersole et al. 2016), what is the impact of self-corrections on a research career? To shed light on these questions, we show results of an ongoing study in which we conducted exploratory interviews with a sample of funding agencies representatives and/or members of review panels from different countries, followed by a survey of reviewers with experience as grant reviewers for US funding agencies.

## Methods

As part of an ongoing project exploring the influence of retractions and self-correcting the literature on the career of scientists, we conducted a qualitative pilot study consisting of six interviews with insight into grant review in several different countries (Ribeiro & Vasconcelos, 2020), followed by a survey of faculty reviewers at one institution. Interviews were conducted with a purposive sample of six expert representatives and/or researchers with experience as review panel members, from the following institutions: one representative of NIH (United States), the major federal agency “conducting and supporting basic, clinical, and translational medical research…” (https://www.nih.gov/news-events/news-releases/nih-nsf-collaborate-accelerate-biomedical-research-innovations-into-marketplace); one researcher from Northwestern University (United States), with experience as a review panel member for NSF (United States), which funds “ research and education in most fields of science and engineering” (https://www.nsf.gov/funding/aboutfunding.jsp); one representative of the European Commission (European Union), the largest funding program in Europe (https://ec.europa.eu/info/research-and-innovation/funding/funding-opportunities/funding-programmes-and-open-calls_en); one representative of the Health Research Board-Ireland (Ireland), the major state funding agency for health research in Ireland (https://www.hrb.ie/about/who-we-are/); one researcher from INSEEC – School of Business & Economics (France), among the largest business schools in Europe (https://www.inseec-bs.com/), with some experience as reviewer for research and non-research proposals; and one representative of São Paulo Research Foundation (FAPESP), the largest state funding agency in Brazil (http://www.fapesp.br/en/6026).

Our interview research protocol was approved by the Research Ethics Committee of the University Hospital of the Federal University of Rio de Janeiro (UFRJ) in April 2017. Interviews were conducted either in person or on Skype. As a follow-up study, we designed a survey instrument with a demographic section and a content section that combined questions from this previous pilot work with additional ones to broaden the scope of the research. The associated research protocol was approved in March 2020, with exempt status, by the Institutional Review Board (Research Ethics Committee) of the University of California San Diego (UCSD), where this follow-up study was conducted. Research participants were research faculty at UCSD, with experience as grant reviewers for NSF, NIH, and/or a few other funding agencies. Online survey invitations were sent to faculty members (n=942) from a wide variety of departments, including the natural sciences, health sciences, engineering, and social sciences. The SurveyMonkey survey was open from October/2020 to February/2021, with three reminder messages in this period. Responses (n= 224, 24 % response rate) were collected for quantitative analysis, and the corpus obtained from the comments of participants excluded only a few sentences that were unrelated to the survey questions or to the issues raised. The associated responses, however, were kept in the dataset. These sentences accounted for less than 0,5% of the whole corpus, and they did not lead to exclusion of the other answers from the same participant.

## Results and Discussion

The main views of representatives of funding agencies and/or researchers with experience as review panel members, from the NIH, NSF, the European Commission, the Health Research Board-Ireland (Ireland), INSEEC Business School, and FAPESP on how they would react to retractions on a CV of an applicant are shown in **Box 1** (Ribeiro & Vasconcelos, 2020). Their comments suggest that major research funders do not have policies in place to address retractions or contributions to the correction of the literature in review criteria. Their comments were shared as personal opinions, based on their experience as grant reviewers, and these opinions do not necessarily reflect those of the institutions these respondents worked for or of the panels they served. However, note that, overall, their comments are consistent with the fact that assessing retractions or self-correction of the literature by an applicant is not part of the review process. As Janke (2018) pointed out, “all grant proposals are evaluated based on universal criteria: quality of the project, achievements of the proposer or consortium, feasibility, and the context of research in this domain”.

### Box 1

Excerpts from participants’ comments regarding the influence retractions may have in the evaluation process of grant applications.

**On the influence of a retraction (s) on a researchers’ CV at the moment that he/she is applying for funding in your institution/agency:**

**Participant 1 - NIH (United States): “**…People make mistakes…but I think many, many mistakes…tend to be more problematic. We don’t have a way right now to track this… we can see all of the papers or articles that are associated with a particular grant, but they don’t always show whether or not they’re retracted or corrected or not. So we don’t have a good way to track that right now. The program official who oversees the grant…hopefully is aware of the field and will pay attention to, you know, when the paper gets corrected or retracted, so we do rely on those individuals as well to be alert. I think it’s important, now especially if there’s an expression of concern or retraction, because it’s possible that the research finding is not true in that case. So then I would be concerned. Why are they… including this in their application if it’s been retracted? I don’t know how we would make sure that someone is including that information [on retractions] and how we would know whether or not…they are truthful about it… Right now we only ask them to include those papers that are relevant to the application, so they may not even need to include those papers if they are not… relevant to the research that they are proposing…. I would be afraid of that. If they included it in the CV…it’s hard to know what the reviewers would think, but my impression is that the reviewers may look at that and wonder why…I do understand that people retract papers for many reasons too and sometimes it’s not due to research misconduct…the word “retraction” I think may be something that we need to think about… not calling it a retraction and calling it something else.

**Participant 2 – Northwestern University (United States):** “…if you’re in a review panel… you know, a major panel for senior faculty… I don’t need to read the CV of people I know. In fact, I probably, I mean, I’ll look at it just to say, “Oh, yeah, I didn’t know they published that.” But… when you’re called to be a reviewer, especially when you reach a particular age in life where you know a lot of people in your field personally, yeah, so the CV is interesting, but for senior people it’s, yeah, it doesn’t much matter. I already know it by looking at who’s proposing… I think the answer to that question is, I don’t know. I don’t know how much data exists on fabricated, falsified, plagiarized material in the literature or in, you know, the preliminary data section of proposals…. I think there it needs to be an analysis of this… if we could have retractions that actually were labeled something other than retraction, just retraction, retraction for, you know, malfeasance, retraction for, you know, materials and methods issues…you pick it. I think we have to advance. But I don’t… So you asked me a question, the question you asked would require me, I think us, as a society, scientific society, to actually understand the depth of the problem…. “.

**Participant 3-INSEEC (France):** “…you have several projects and then if for every project you have things that are being retracted it might be stopped, but, again, it’s not considered individually. I mean, in a CV… usually CVs are built, so in a CV you might not see that there is a retraction and you might not see, and if you see it… because retraction cannot be best practices. You can have cases of people that are not English speakers and you might consider that it’s not properly written and retract it. I believe I’ve seen this in retraction quite a lot. So it happens. And then what I also believe is that retraction only happens after the first. So I don’t think retraction is something, when you have a fraud case, a potential fraud case, I believe that you will have a retraction, volunteer retraction, for people that are really in best practices and really want to amend things and make things better. … But for me it’s not, I mean, it’s like the burden of evidence or the burden of proof; you need to have several issues to really impact a review. I don’t really, I don’t know…again, it’s something very… It’s only an opinion, but I really don’t think that this might be paramount to the rest of the evaluation. What is important, again and again, is this going to put the project at risk or not?”.

**Participant 4-European Commission (European Union)**: “… both retractions and corrections are extremely important for the scientific record. They have major importance… Retractions can happen, even (with) no misconduct…. We set definitions of misconduct, it’s an opposite, different story. So both of them are important for getting the scientific record straight… How do you…apply the same… evaluation criteria on retractions in Mathematics, Astronomy or Social Sciences? We do not have the experience internationally on these situations, not only in the European Commission… nobody does. That idea would be a very good idea, but it will probably lead to a very steady procedure and, you know, I mean, thinking about it, not spend time in actually designing the procedure, but how retractions can be used. I can assume that the beginning will be very heavy until the system goes through the growing pains and learns how to deal with it. If it’s a good idea, I would prefer that instead of having a perfect CV, remember about that movement that started in the US, I think it started at Princeton, we had the failure CV. Not only how many successes I had, how many grants I got, how many papers I published, but also how many grants I didn’t get, how many papers I didn’t publish, because there was that, so what it was called the “failure CV”“.

**Participant 5 – Health Research Board (Ireland)**: “Well, they have no way of knowing whether a paper has been retracted if they are not told about it, unless where this might actually come into play is if it’s a senior fellow asking, where you’re… planning to give them a very large amount of money. In that case, we probably would ask them for all of their publications. And if they’re seniors, they’re probably going to be known to the panel, the peer review panel anyway. So if there are issues around the publications, if they have had difficulties around retractions…it’s a very small system; that (will) probably… be known. But we don’t systematically check every publication or check that person’s name, let’s say, in PubMed, to see whether or not they’re associated with retracted publications. So that’s not the information we chase; they can choose to tell us about those retractions if they wish, but I really haven’t seen that happen…Well, I think it would depend on the reason for the retraction. So if the retraction was because they found a flaw in the publication and they retracted it in an attempt to correct, there’s absolutely no issue with that…. If, on the other hand, we discovered that somebody had had one or more publications retracted due to misconduct, then I think that that would probably and prompt us to either not fund them and… it would depend on when this happened, of course. I mean, if it’s a very recent thing, then, you know, the chances are we might not fund them, because a) they haven’t told us about this, so they’re being dishonest in how they’ve presented themselves…but it’s a very hard thing for someone to tell a funding agency about…. If they don’t tell us about it and we find out, well, then they seem to be dishonest, but if they do tell us about a misconduct case or hear a finding that they have against them, then… I think it would definitely bias the way the peer review panel would look at them”.

**Participant 6 - FAPESP (Brazil):** “…the number of retractions is very small, so, it will take time to affect the evaluation system, I think… because few researchers have retractions and even fewer have a significant number of retractions and [there] are still researchers with a solid background of research that remain active… you asked if in the near future there will be some influence… I do not see much reason to, but the way things are in the world and in Brazil, moralism and the accusations of enemies…, probably this will enter the agenda…my perception is not that it will take a long time… My answer is that I do not think it would be necessary, but I think it will happen because this whole subject is dominated by a moralism and a political correctness that is beyond rationality, because it is a matter generally brought to allegations and persecutions. The great difficulty that I see in the rational treatment of this problem is that, in many cases, the reason for the retraction or the claim of…non-reproducible results or other things - has a great tendency to become an instrument of personal revenge between people… Scientists know that the fact that an article that is published does not mean that it is right ….every scientist who reads the article has to make their analysis, their criticism… Often the person carries out their own work and says, ’Unlike the results obtained by so-and-so, here I got this because in my case I did better or because it is a different situation’ … It turns out that after the scientific literature went to the internet, and it has become something that not only scientists read, it has become a different thing, because the lay public and journalists assume that if it came out in the scientific literature, it is right and the responsibility for being right is of the editor of the journal and the reviewers. And then it became something totally different…”.

As can be seen in **Box 1**, these respondents demonstrate that review criteria in their institutions and/or on the panels where they served in the capacity of reviewers do not encompass discussions of retractions or self-corrections of the literature in the assessment of applications.

Participant 4 (European Commission), for example, says that

> *“We do not have the experience internationally on these situations, not only in the European Commission… nobody does”*.

The following statement by Participant 1 (NIH) is aligned with that by Participant 4. This respondent 1 comments that

> *“We don’t have a way right now to track this… we can see all of the papers or articles that are associated with a particular grant, but they don’t always show whether or not they’re retracted…. or corrected or not. So we don’t have a good way to track that right now*.*”*

For Participant 2, with experience at reviewing for NSF, the perception is similar:

> *“…if you’re in a review panel… you know, a major panel for senior faculty… I don’t need to read the CV of people I know. In fact, I probably, I mean, I’ll look at it just to say, “Oh, yeah, I didn’t know they published that*.*” But… when you’re called to be a reviewer, especially when you reach a particular age in life where you know a lot of people in your field personally, yeah, so the CV is interesting, but for senior people it’s, yeah, it doesn’t much matter. I already know it by looking at who’s proposing… I think the answer to that question is, I don’t know”*.

Participant 5 (Health Research Board, Ireland) comments that

> *“…we don’t systematically check every publication or check that person’s name, let’s say, in PubMed, to see whether or not they’re associated with retracted publications. So that’s not the information we chase; they can choose to tell us about those retractions if they wish, but I really haven’t seen that happen…Well, I think it would depend on the reason for the retraction. So if the retraction was because they found a flaw in the publication and they retracted it in an attempt to correct, there’s absolutely no issue with that…*.*”*.

Although this is a small group of respondents and does not offer definitive answers, the review panels where they served were for agencies with a strong role in the international panorama of research funding, which makes these views particularly relevant. All of them were aware of the difficulty of dealing with corrections of the literature, particularly retractions, in the context of funding decisions:

> *I believe that you will have a retraction, volunteer retraction, for people that are really in best practices and really want to amend things and make things better. … But for me it’s not, I mean, it’s like the burden of evidence or the burden of proof; you need to have several issues to really impact a review. I don’t really, I don’t know…again, it’s something very… It’s only an opinion, but I really don’t think that this might be paramount to the rest of the evaluation. What is important, again and again, is this going to put the project at risk or not?”*.

Participant 3 (INSEEC, France).

In a nutshell, these respondents seem to recognize that funding bodies will have to deal with the challenge of corrections to the research record soon and for different reasons. For Participant 6, dealing with retractions

> *“…would be necessary, but I think it will happen because this whole subject is dominated by a moralism and a political correctness that is beyond rationality, because it is a matter generally brought to allegations and persecutions. The great difficulty that I see in the rational treatment of this problem is that, in many cases, the reason for the retraction or the claim of…non-reproducible results or other things - has a great tendency to become an instrument of personal revenge between people… Scientists know that the fact that an article that is published does not mean that it is right”*.

As reflected in these respondents’ views, retractions and the whole dynamics of correcting the literature, are a recent phenomenon. Doubts abound among their comments. Those from the NIH and European Commission, for example, illustrate this factor. Doubts also reflect the apparent lack of formal policies to guide review panel members concerning applications by PIs that had experienced retractions. Another factor is the one raised by Participant 6 (from FAPESP), who called attention to bad incentives involved in retractions driven by misconduct allegations and to demands associated with retractions created by the higher public exposure of science. Worth noting is that the doubts expressed in these conversations are from funders/review panel members at major institutions in countries with a considerable share of publications (Scimago Journal & Country Rank 2019). Also, whereas the results offer only a partial picture of the problem, they resonate with an international debate over the reward systems of science (Casadevall 2019). This debate involves critical attitudes toward traditional views of credit and impact in science (Curry et al. 2020), which invites academia to have a broader look at retractions and correction of the literature at large.

### Perceptions about retractions, among grant reviewers in the US, and grant review criteria

Retractions of publications is a quite recent phenomenon in academia, as can be seen in the results from the pilot survey of research faculty at UCSD with experience as reviewers for major funding agencies in the US. The respondents were predominantly male (70%), full professors (71%), and in the natural sciences and engineering (70%). In terms of experience as grant reviewer, most (52%) reported more than 20 years.

Overall, respondents of the survey, who declared having served on review panels for NSF (80%) and/or NIH (40%), indicated they almost never come across retractions in publications in their fields let alone in the material included in grant applications. In fact, their responses, harmonized with those shown in **Box 1**, indicate that looking at retractions and, more broadly, on self-corrections of the literature among applicants, does not have any role in the grant review process. This finding is consistent with responses to the question on rating the importance these research participants would give to a list of given items when reviewing a grant proposal, as shown in **Figure 1**:

**Figure 1.**
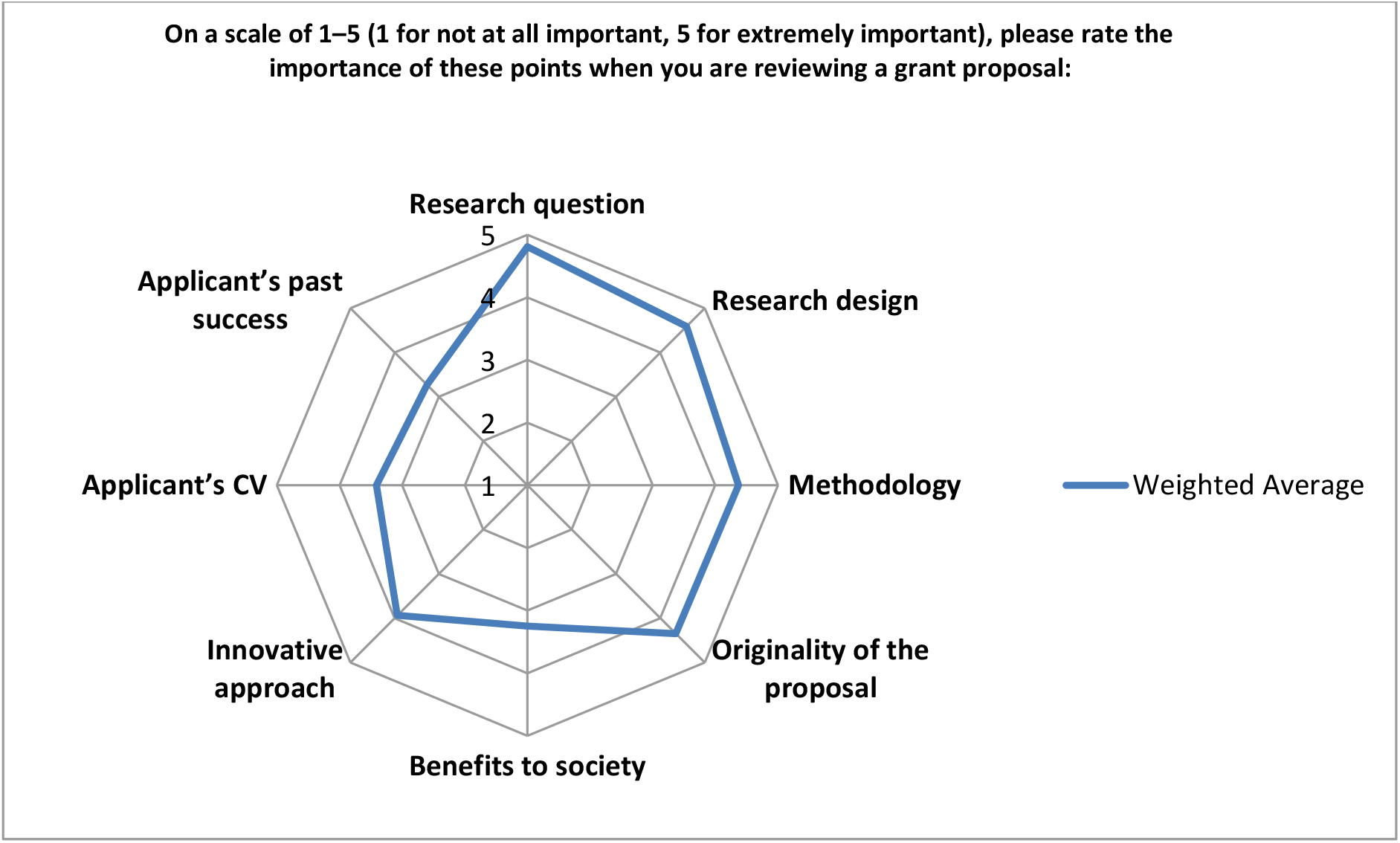
Items rated by respondents (n=185) according to the importance they give to each while reviewing a grant proposal.

These results are to be expected given that review criteria usually focus on research design, novelty, innovation, impact, and the research team. Falk-Krzesinski and Tobin (2015) identified “eight key questions considered by reviewers of research grant proposals and the associated review criteria terms used by 10 US federal funding agencies”. These authors compared research grant proposal review criteria used by these 10 agencies and found that they are “remarkably aligned” and that “there exist only a small and finite number of ways that federal research grant proposals are actually evaluated”.

### A changing culture of assessment – much room for broader looks at good research practices and the correction of the literature

This culture of assessment of grant proposal does not take into consideration good research practices that can strengthen the reliability of the research record. In fact, the role of self-retractions, for example, for honest errors or any problem that may compromise the validity of published results is not captured by the rationale for most grant review criteria.

However, in our changing landscape for the responsible assessment of research and researchers, funding agencies have started to take a proactive role in discussing grant review criteria. In the last few years, funders across the globe have revisited several aspects of review criteria, and discussions at a recent Global Research Council (GCR) conference, which included contributions from the NSF and other leading agencies, is helping to reshape the culture of assessment (GRC 2020). In January 2022, Hatch and Fritch (2022) reported that “many funding organizations are shifting to a more holistic interpretation of research outputs and achievements that can be recognized in the grant evaluation process through the use of narrative CV formats that change what is visible and valued within the research ecosystem. As communities of practice in research and innovation funding, the San Francisco Declaration on Research Assessment (DORA) funder’s group and Funding Organisations for Gender Equality Community of Practice (FORGEN CoP) partnered to organize a workshop focused on optimizing the use of narrative CVs, which is an emerging method used by research funders to widen the research outputs that can be recognized in research assessment.” This initiative demonstrates that there is much room for improvement not only in terms of research outputs but also in terms of attitudes toward the evaluation of an applicant’s actual contributions to academia.

As the comments of the six respondents in our pilot study suggest, there is plenty of room for the research community to address the role correcting the literature can or should play in funding decisions. Consistent with this argument, the 2017 revised code of conduct of All European Academies acknowledges that giving authors “credit for issuing prompt corrections post publication” (ALLEA 2017) is among best practices in the scientific enterprise. As to why retractions are not normally considered in the grant review process, nearly 60% of respondents reported the perception that retractions are a “rare phenomenon”, and 30% of respondents indicated that it is too early to assess the role retractions should have in the review process (**Figure 2**).

**Figure 2.**
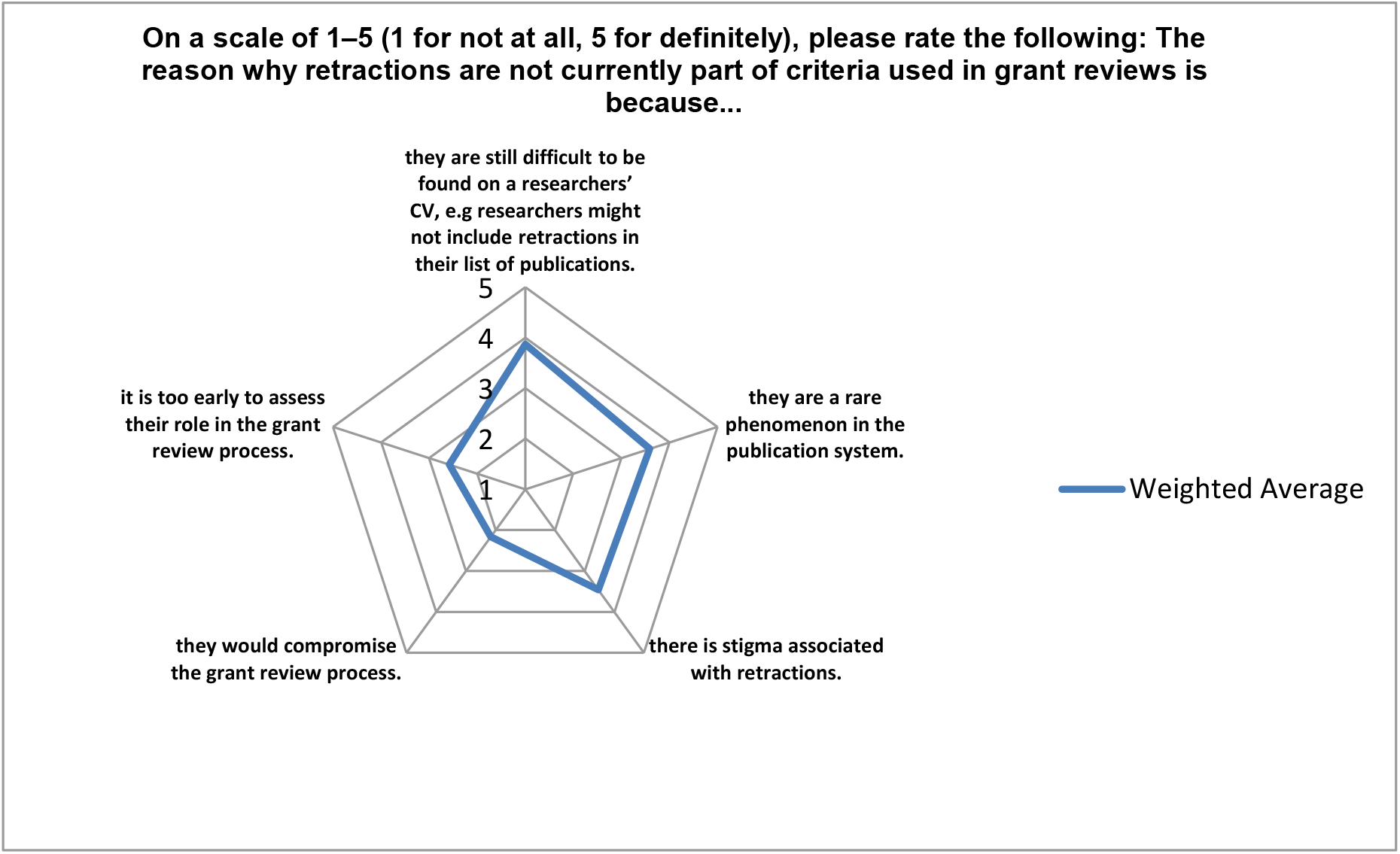
Patterns of response (n=170) for options offered to respondents to explain why retractions are not currently part of criteria used in grant review.

This attitude reflects transformations that the reward systems of science are undergoing, particularly in the biomedical sciences. Reporting on progress and priorities for the Reproducibility2020 Initiative, launched in 2016 by GBSI (Global Biological Standards Institute), Freedman et al. (2017) mentioned some of the actions. Among these is to “explore new incentive structures for career advancement that move away from the traditional impact factor and funding paradigms to reward greater data and methods transparency, adherence to best practices and standards…”. This provocative call for changing “funding paradigms” is consistent with the discussion of weaknesses in criteria used in peer review processes for grant applications (Mutz et al. 2012; Bendiscioli 2019). Discussing some of the weaknesses, Bendiscioli (2019) claims that “funders have the responsibility to ensure that their money is being distributed in the most efficient and fairest way and should not shy away from experimenting with new ideas to achieve this goal”. As Casadevall (2019) notes for the culture of rewards in the biomedical sciences, there must be a change and “… review and promotion committees must take a different approach in evaluations.”

Retractions due to self-correcting the literature are not yet incorporated into the scientific culture, but repercussions are still evolving in the context of research rewards and assessment. This fact is illustrated in the comments of the six respondents in **Box 1** and in those of 32 of the 86 survey respondents who added comments following the question in which they were asked to rate the importance of “…factors influencing your judgment in grant review, when it comes to the CV of the Principal Investigator (PI)” (n=184). (**Box 2**) As can be seen in **Figure 3**, one of the options was “Retractions or other types of corrections in the PI’s publication record”.

**Figure 3.**
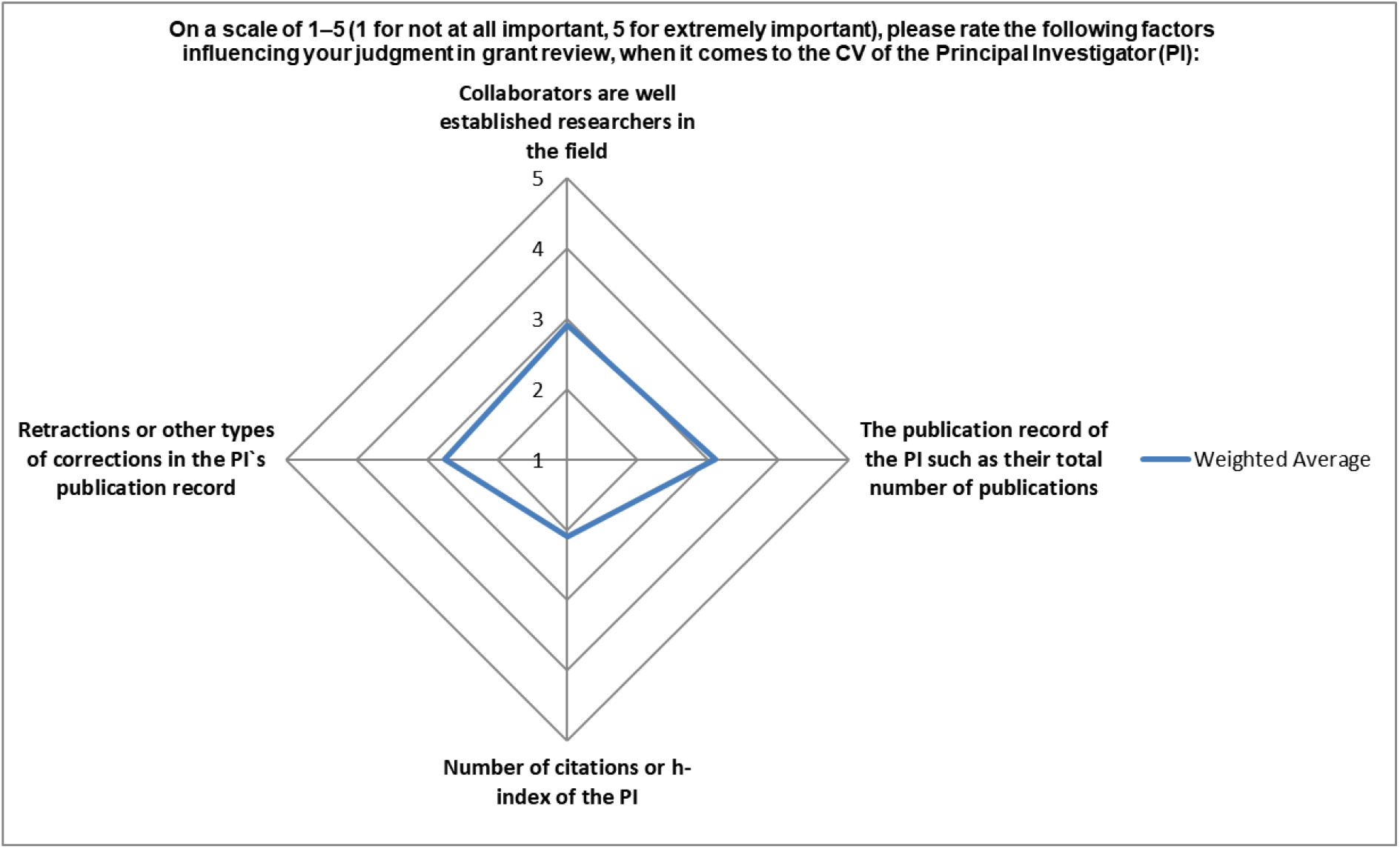
Patterns of response (n=184) for options offered to respondents to rate factors influencing their judgment in grant review, considering the CV of the Principal Investigator (PI).

#### Box 2

Comments, separated by a slash, from 32 reviewers for US funding agencies out of 86 that added comments to the question “On a scale of 1–5 (1 for not at all important, 5 for extremely important), please rate the following factors influencing your judgment in grant review, when it comes to the CV of the Principal Investigator (PI)” (n=184).

**1 Unless you look for it you won’t readily be aware of retractions when reviewing grants. / 3 I have never had to evaluate PI with a record of retractions. / 6 I have never reviewed a grant where the PI had a retraction, only ‘correction’ / 7 I have never seen a retraction or correction to the record, so it is hard for me to say much about this issue/ 11 Never had an application with retraction to deal with / 13 I have reviewed dozens of proposals but I don’t recall ever considering or weighing retractions. / 15 I actually don’t recall *ever* seeing a grant application that mentioned a retraction. / 17 I don’t remember seeing retractions / 20 Never thought about retractions. Not sure how common they are in economics though. / 23 I have not seen instances of retractions / 25 [W]hen reviewing for the NSF I only see the biosketch and it has never contained retractions as far as I’ve seen. In the biosketch you only get to include 5 products related to the research, so there is rarely room to be comprehensive. / 28 I have encountered very few (perhaps 0) such situations. Retractions are rare and I may even have missed them. / 29 I have never seen a retraction on a CV / 31 This information is not provided and I wouldn’t know how to get it / 35…how much it would affect would depend on why the retraction happened. If it happened due to an error, it would not weigh heavily in my view. If it was cheating, then it would be very important and I doubt there’s anything else that would make me be positive in the proposal. / 37 I have never seen a retraction on a PI’s publication record, but it would give me pause if I did. I’d want to know more about how it happened/ 38 [N]ever looked for retractions, that didn’t cross my mind / 42 [N]ever encountered a proposal that included such corrections / 45 I’ve never come across a PI with retractions as a reviewer / 46 I have never seen any retractions / 48 I have never seen a retraction listed on a CV / 57 I’ve never seen a proposal in which the PI had a non-negligible correction or retraction. / 58 I’ve never had an experience where I knew of any retractions. / / 62 I have never actually seen retractions in a CV. These are rare, and unless explicitly required, I would expect them to be omitted. CVs are essentially 100% positive. /66 [N]ever seen one. can’t comment. /67 I have no memory reviewing a proposal from a PI with a retraction. / 73 I’ve not had an applicant with retractions / 74 I have never seen a retraction involving a grant proposal / 75 I’ve never seen a retraction. Correction I’ve seen and I don’t think they’re typically a big deal (actually I thin[k] it predisposes me towards the PI since it shows that they care about integrity and want to make sure that the most correct version of their work is out there) /80 I never checked the retraction/corrections by the PIs during proposal review / 81 I’ve never seen a retraction on a CV/ 84 I have not really encountered major retractions or corrections**.

Despite the comments in **Box 2**, which corroborate the fact that retractions are rare, not easily tracked in the grant review process, and not part of grant review considerations, the majority of respondents recognized that mechanisms for correcting the scientific literature are a way to strengthen the reliability of the research record, with 90% rating it at 4 (24%) or 5 (66%) (**Figure 4**).

**Figure 4.**
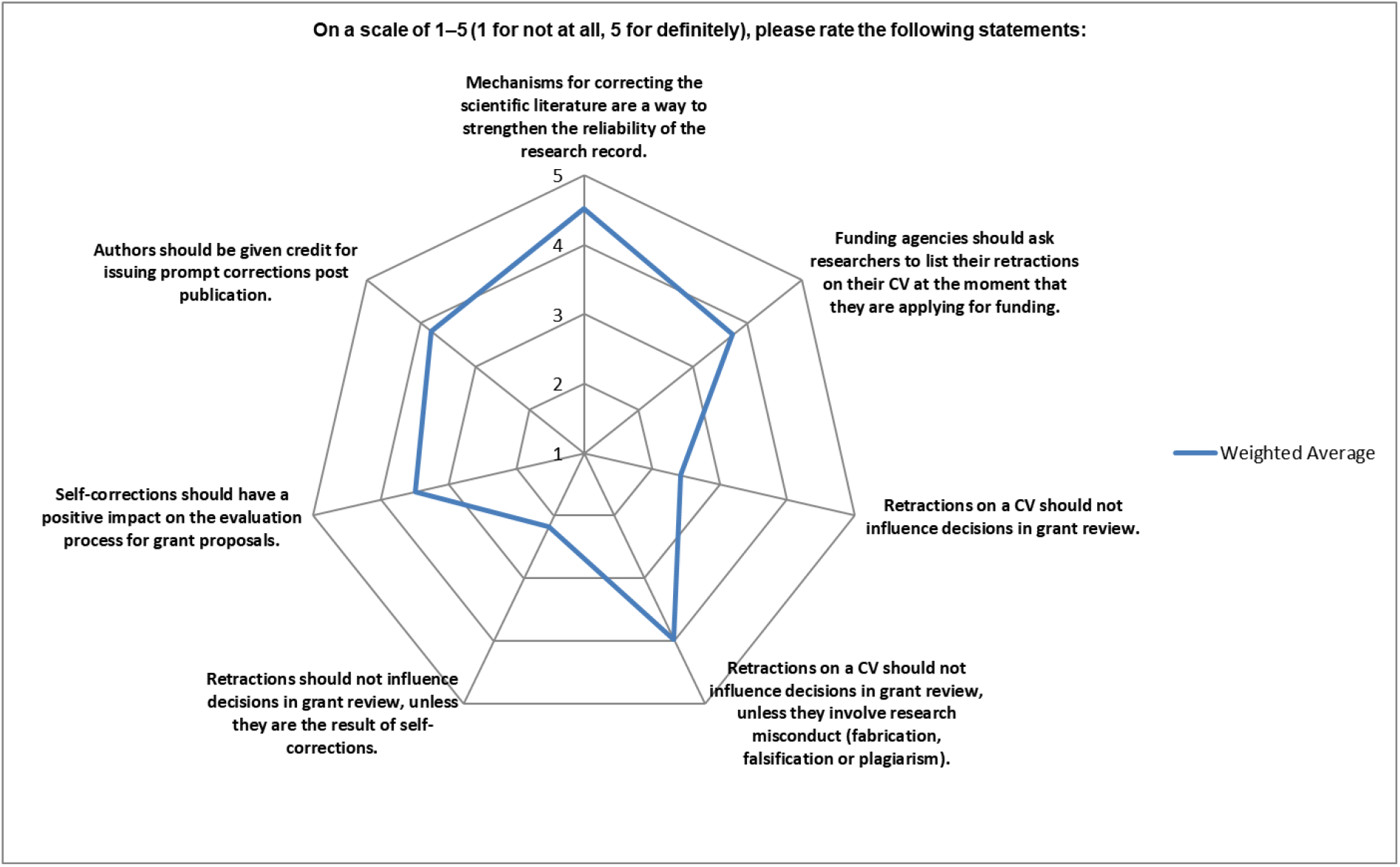
Patterns of response (n=169) for statements about retractions, self-correcting the literature and their role in decisions in grant review.

This recognition among the greatest fraction of respondents echoes what other researchers have declared about the role the correction of the literature plays to strengthen science and public trust in the research community. Yet, when it comes to funding criteria, only a few actions, such as that by ALLEA (2017) have acknowledged that correcting the research record should be recognized as good research practice. As an editorial in Nature Human Behavior (2021) has recently pointed out, “[r]etractions are a key tool for maintaining the integrity of the published record. We need to recognize and reward researchers, especially early-career researchers, who do the right thing in coming forward with a request to retract research that cannot be relied upon due to honest error.”.

Yet, there are certainly several shades of gray in the approach to retractions and what they mean or should mean in the career of researchers, especially because most retractions have been issued for research misconduct and not for honest errors or other “positive” reasons (Fang et al. 2012; Ribeiro & Vasconcelos 2018; Wang et al. 2019). As we have previously mentioned, this type of correction in a researcher’s publication record may have detrimental effects on her/his career in terms of citation patterns (Furman et al. 2012; Lu et al. 2013), productivity and even debarment for funding in some extreme cases (Mongeon and Larivière 2016; Azoulay et al. 2017). Overall, these consequences reflect on the relevance more of retractions for misconduct than those for honest error. In the context of funding, retraction reasons, including those related to honest error should not be overlooked in grant review, especially now that notions of rewards and allocation of credit in science have been revisited and gained renewed attention.

Especially in the last two decades, correcting the literature as a mechanism to improve science has added another layer of complexity to assess and validate scientific knowledge. Yet, as highlighted in **Figure 4**, this new research landscape, which brings an even sharper focus on “the credibility of the published literature” (Soderberg et al. 2021), is not reflected in funding criteria. That said, it is essential to broaden our view of retractions in the research endeavor and make self-correction of the literature part of the reward systems of science (Fanelli, 2016). Different actors in academia have been calling for changes in the current science model to *collectively* improve the way research is funded, done, reported, reviewed, and communicated to peers and to the public.

## Conclusion

The results from our pilot project and the follow up survey among US grant reviewers show that the majority of participants recognize that mechanisms for correcting the scientific literature are a way to strengthen the reliability of the research record. However, this finding does not lead to expectations for including retractions, for misconduct or honest error, in the grant review process. Our results also show that the research question, research design, methodology, and originality of the proposal, and not the applicant’s CV, are decisive for grant decisions.

When it comes to retractions by an applicant in the grant review process, as they are not usually listed on CVs, their influence on funding seems null up to this point. Wider discussion and consideration of the role this process of correcting the literature should have in the research system is timely, especially considering that the whole point of correcting the research record is to strengthen science at large.

## Acknowledgments

We thank the grant reviewers and representatives of funding agencies who contributed to this study. CAPES is also acknowledged for the support granted to the first author (Ribeiro, MD) to conduct a Sandwich PhD period at UCSD.

## References

All European Academies (ALLEA). (2017). The European Code of Conduct for Research Integrity. Available at: https://ec.europa.eu/research/participants/data/ref/h2020/other/hi/h2020-ethics_code-of-conduct_en.pdf

Azoulay, P., Bonatti, A., Krieger, J. (2017). The Career Effects of Scandal: Evidence from Scientific Retractions. Research Policy, 46(9): 1552–1569

Bendiscioli, S. (2019). The Troubles with Peer Review for Allocating Research Funding. EMBO Reports, 20 (12): e49472. https://doi.org/10.15252/embr.201949472

Casadevall, A. (2019). Duke University’s huge misconduct fine is a reminder to reward rigour. Nature, 568 (7). https://doi.org/10.1038/d41586-019-01032-w

Curry, S., de Rijcke, S., Hatch, A., Pillay, D.G., van der Weijden, I., Wilsdon, J. (2020). The changing role of funders in responsible research assessment: progress, obstacles and the way ahead. Research on Research Institute. Report. https://doi.org/10.6084/m9.figshare.13227914.v1

Deutsche Forschungsgemeinschaft (DFG). (2018). Rules of Procedure for Dealing with Scientific Misconduct. Available at: https://www.dfg.de/formulare/80_01/80_01_en.pdf

Ebersole, C.R., Axt, J.R., Nosek, B.A. (2016). Scientists’ Reputations Are Based on Getting It Right, Not Being Right. PLOS Biology. https://doi.org/10.1371/journal.pbio.1002460

Falk-Krzesinski, H.J., Tobin, S.C. (2015). How Do I Review Thee? Let Me Count the Ways: A Comparison of Research Grant Proposal Review Criteria Across US Federal Funding Agencies. The Journal of Research Administration, 46(2):79–94.

Fanelli, D. (2016). Set up a ‘self-retraction’ system for honest errors. Nature, 531: 415. https://doi.org/10.1038/531415a

Fang, F.C., Steen, R.G., Casadevall, A (2012). Misconduct accounts for the majority of retracted scientific publications. Proceedings of the National Academy of Sciences, 109(42):17028–33. https://doi.org/10.1073/pnas.1212247109.

Global Research Council (GRC). (2020). Responsible Research Assessment. Available at: https://www.globalresearchcouncil.org/news/responsible-research-assessment/

Hatch, A., Fritch, R. (2022). Cross-funder action to improve the assessment of researchers for grant funding. Available at: https://sfdora.org/2022/01/19/cross-funder-action-to-improve-the-assessment-of-researchers-for-grant-funding/

Janke, C. (2018). A unified reviewing format for grant applications and evaluations. EMBO Reports, 19 (2). https://doi.org/10.15252/embr.201745611

São Paulo Research Foundation (FAFESP) (2012). Code for Good Scientific Practices. Available at: http://fapesp.br/boaspraticas/FAPESP-Code_of_Good_Scientific_Practice_jun2012.pdf

Freedman, L. P., Venugopalan, G., & Wisman, R. (2017). Reproducibility2020: Progress and priorities. F1000Research, 6, 604. https://doi.org/10.12688/f1000research.11334.1.

Furman, J.L., Jensen, K., Murray, F. (2012). Governing Knowledge in the Scientific Community: Exploring the Role of Retractions in Biomedicine. Research Policy, 41(2): 276–290.

Galbraith, K.L. (2017). Life After Research Misconduct: Punishments and the Pursuit of Second Chances. Journal of Empirical Research on Human Research Ethics, 12 (1): 26–32.

Lu, S.F., Jin, G.Z., Uzzi, B., Jones B. (2013) The Retraction Penalty: Evidence from the Web of Science. Science Reports. https://doi.org/10.1038/srep03146

Mongeon, P., Larivière V. (2016) Costly Collaborations: The Impact of Scientific Fraud on Co-Authors’ Careers. Journal of the Association for Information Science and Technology, 67(3): 535–542.

Mutz, R., Bornmann, L., Daniel, H-D. (2012). Heterogeneity of Inter-Rater Reliabilities of Grant Peer Reviews and Its Determinants: A General Estimating Equations Approach. PLoS ONE, 7(10): e48509. https://doi.org/10.1371/journal.pone.0048509

Nature Human Behavior 5: 1591 (2021). https://doi.org/10.1038/s41562-021-01266-7.

Ribeiro, M.D., Vasconcelos, S.M.R. (2018). Retractions Covered by Retraction Watch in the 2013-2015 period: prevalence for the most productive countries. Scientometrics, 114(2): 719–734. https://doi.org/10.1007/s11192-017-2621-6

Ribeiro, M.D., & Vasconcelos, S. (2020, January 10). Should corrections of the literature influence grant review?. https://doi.org/10.31234/osf.io/49vpa

Scimago Journal & Country Rank (SJR). (2019). Available at: https://www.scimagojr.com/countryrank.php

Soderberg, C.K., Errington, T.M., Schiavone, S.R. et al. Initial evidence of research quality of registered reports compared with the standard publishing model. (2021). Nature Human Behavior 5, 990–997. https://doi.org/10.1038/s41562-021-01142-4

Wang, T., King, Q.R., Wang, H., & Chen, W. (2019). Retracted Publications in the Biomedical Literature from Open Access Journals. Science and Engineering Ethics, 25: 855–868. https://doi.org/10.1007/s11948-018-0040-6

